# Analysis of individual identification and age-class classification of wild female macaque vocalizations without pitch- and formant-based acoustic parameter measurements

**DOI:** 10.1101/2025.03.17.643698

**Authors:** Rentaro Kimpara, Fumiya Kakuta, Hiroki Koda, Ikki Matsuda, Goro Hanya

**Author notes:** Correspondence to RK.

## Abstract

In recent years, deep learning has achieved high performance in bioacoustic classification tasks by leveraging automatically processed acoustic features for large datasets. However, few performance evaluations of automatically processed acoustic features have been conducted on small-scale data because deep learning requires large datasets. To test whether mel spectrograms (an automatically processed acoustic features) are effective for classifying relatively small acoustic data, we evaluated the performance of two classification machines (random forest and support vector machine) using mel-spectrograms of 651 coo calls of six wild female Japanese macaques on two tasks: 1) individual identification and 2) age-class classification between younger (<10 yrs) and the older animals (>20 yrs). For the individual identification task, the mean balanced accuracy was 0.81 for random forest and 0.82 for support vector machine. For the age-class classification task, the mean balanced accuracy was 0.91 for random forest and 0.93 for support vector machine. Considering that of all the calls were recorded in the wild, methods using automatically processed acoustic features, such as mel spectrogram, are effective in classifying small acoustic data for the individual identification task. The high performance in the age-class classification task might be attributable to the potential of mel spectrograms to capture the characteristics of older individuals (e.g. harshness).

## Introduction

To elucidate the vocal communication of animals, bioacoustic studies have implemented tasks to classify calls from acoustic features for such purposes as identifying individuals making vocalizations and classifying vocal repertoires. This work has been pursued for more than half a century (Arnaud et al., 2023; Fukushima et al., 2015; Marler, 1956; Marler and Hobbett, 1975; Mitani, 1986; Rendall et al., 1998; Winter et al., 1966). Historically, conventional acoustic analysis has measured numerous acoustic features like pitch and formant and has examined classification performance using dimensionality reduction and multivariate analysis (Ceugniet and Izumi, 2004; Oyakawa et al., 2007; Rendall et al., 1998). Reported classification accuracies were 83.7-92.3%, 90.5-98.4%, and 63.6-90.9%, respectively. Although these conventional approaches are still in use, they have suffered from problems such as the arbitrary nature of selecting acoustic features and the lack of versatility due to the definition of species-specific acoustic features.

Recent machine learning, especially deep learning, can automatically process acoustic signals for classification tasks without explicitly measuring specific features, which solves the problems associated with the conventional method (Arnaud et al., 2023; Lakdari et al., 2024). Nevertheless, current deep-learning techniques have a methodological shortcoming in that they typically require a large amount of supervised data, often ranging from thousands to millions of annotated samples in animal sound classification tasks (Nanni et al. 2020; Heinrich et al. 2025). This is an obstacle to building models because preparing a large amount of supervised data requires a huge amount of time or is nearly impossible due to difficulties such as collecting sufficient data in the field. Therefore, developing classification models for relatively small amounts of data without measuring specific acoustic features could alleviate the difficulty of model building and facilitate bioacoustic studies of many species. However, no such models have so far been built.

Coo call is a contact call of macaques and accounts for a large part of Japanese macaque vocalizations (Koda and Sugiura, 2010). Acoustically, the characteristic of coo call is a tonal sound, that is to say, the fundamental frequency of the harmonics is clear (Green, 1975). Concerning individual identification from coo calls of Japanese macaques, Ceugniet and Izumi suggest that call duration and start and end fundamental frequency are crucial (Ceugniet and Izumi, 2004), and Mitani pointed out that fundamental frequency is less variable in the same individual compared to call duration (Mitani, 1986). Likewise, formant frequencies—which are generally determined by vocal tract length and shape—have been examined to describe acoustic properties relevant to vocal identity. A study in rhesus macaques, a closely related species, demonstrated that vocal tract characteristics contribute to individual discrimination (Rendall et al., 1998), and this finding has been supported by experimental work in Japanese macaques (Furuyama et al., 2016). These studies attempted to identify individuals based on vocalizations by measuring specific acoustic characteristics. However, to date, no research has applied automatically extracted acoustic features to distinguish Japanese macaques based on their vocalizations.

Acoustic features are relatively unstable due to age-related changes. Aging in animals is associated with physical and cognitive changes (Hindle et al., 2009; Manrique and Call, 2015). In humans, aging physio-morphologically affects acoustic features such as fundamental frequency and signal-to-noise ratio (Abitbol et al., 1999; Ferrand, 2002; Pontes et al., 2006; Ryan and Burk, 1974; Stathopoulos et al., 2011), suggesting that the age of a human could be identified from the voice (Harnsberger et al., 2008; Huntley et al., 1987). Interestingly, some studies suggest that impressions of the speaker are influenced by age-related acoustic features such as pitch (Feinberg et al., 2005; Klofstad et al., 2015; Tigue et al., 2012). While fewer studies have explored senescence-related acoustic changes in non-human animals, some research suggests that such changes influence the listener’s perceptual estimation of caller age based on vocalization (Lemasson et al., 2010; Zipple et al., 2020). However, bioacoustics research has largely focused on early life stages, with limited attention to senescence (reviewed by Ey et al. 2007).

Individual identification of vocalizations or classification of vocal repertoires, notably in group-living primates, has been attempted repeatedly over the past half-century (Arnaud et al., 2023; Fukushima et al., 2015; Marler and Hobbett, 1975; Mitani, 1986; Rendall et al., 1998). However, few studies have attempted to identify vocalizers or classify age classes in wild primates without explicitly measuring specific acoustic features such as pitch or resonance frequency. In this study, therefore, we address the tasks of vocalizer identification and age-class classification (younger: <10 yrs or older: 20 > yrs) in wild Japanese macaques (*Macaca fuscata yakui*) without measuring specific acoustic features.

## Materials and methods

### Recordings of vocalization

The Japanese macaques (*Macaca fuscata yakui*) of Yakushima Island in southern Japan (30.41N, 130.41E), have been studied intensively since the 1970s, and multiple groups have been identified (Yamagiwa and Hill, 1998). We investigated one of the identified groups, named the Petit group, from October to December 2023. At that time, the Petit Group was composed of about 33 individuals, including 10 adult females (>=6 years), 5 or more adult males (>=6 years), 8 juvenile females (1–5 years), 7 juvenile males (1–5 years), and 3 infants. We selected 10 adult females as focal animals and collected vocalization data using the focal animal sampling method, where RK and FK followed the focal animal. The reason we only focused on adult females is that we concerned that male individuals vocalize infrequently (Mitani, 1986), making it difficult to obtain sufficient vocalizations in the wild. One observation session lasted two hours, and we randomly changed focal individuals from those with the least observation time. We maintained a distance of 3 meters or less from the focal individual. We did not follow estrous females although the observation period included mating season. We recorded vocalizations with annotations of the vocalizer using an audio recorder with two channels and 32-bit-float/48 kHz sampling rate formats (48 kHz), which was connected to a directional microphone (Sennheiser MKE600) for recording the animal’s vocalizations and a lavalier microphone (COMICA CVM-V02O) for recording the observer’s annotations, such as recording vocalizers. The total observation time was 112 hours over 20 days. We created classifiers for six adult females (Kapa, 9 yrs; Rine, 8 yrs; Sasa, 7 yrs; Sazae, estimated 21-22 yrs; Taiko, estimated 23-25 yrs; Taiwu, 9 yrs), for which we obtained sufficient high-quality vocal data. The other four adult females were excluded because they had fewer than 80 high-quality calls. The genetic relationship of the six adult females used for machine learning is not fully known, but two pairs (Sazae and Sasa; Taiko and Taiwu) are mother-daughter pairs.

### Analysis

#### Data selection and preprocessing

We used the acoustic analysis software Praat (Boersma, 2024) for annotation to select data in the recordings for which we could reliably identify the vocalizer. After the annotation, we selected the calls that met all five of the following criteria for further analysis: 1) Little background noise, 2) coo call (see Koda and Sugiura, 2010 for its common definition and function), 3) no overlap with other individuals, and 4) certainty about the vocalizer, and 5) less than 1 second, based on the information by Mitani (1986) and Sugiura (1998). From 1181 audio recordings, we finally selected 116 calls for Kapa (102 calls excluded), 89 for Rine (149 calls excluded), 138 for Sasa (84 calls excluded), 122 for Sazae (84 calls excluded), 84 for Taiko (77 calls excluded), and 102 for Taiwu (34 calls excluded), and we used them in subsequent models for classification tasks. Noted that even though the recordings were made continuously and the time of recording is the same among individuals, thereby minimizing bias, it cannot be denied that bias may have been introduced during the data preprocessing. Situations prone to noise, such as those with strong winds and proximity to streams, probably tended to be removed.

#### Generation of Mel Spectrograms

A mel spectrogram is a set of acoustic features derived through logarithmic scale filters designed using the frequency-sensitivity characteristics of human auditory perception, (Stevens et al., 1937) and it is often used values in machine learning (Baowaly et al., 2024; Dossou and Gbenou, 2021; Walsh et al., 2023; Xie et al., 2024). We also used mel-spectrograms in this study, as previous work showed that UMAP successfully classified animal call types, such as those of meerkats (Thomas et al., 2022).

We generated mel spectrograms of the vocal region with Python’s audio processing module *librosa* (McFee et al., 2015). The original code for the generation of mel spectrograms was based on a previous bioacoustics study on meerkat calls including seven call types (Thomas et al., 2022), and we set the key parameters of generating mel spectrograms generations as follows: N_MELS=40 (number of Mel bands), FFT_WIN=0.03 (30-ms window size for STFT), FFT_HOP=0.00375 (hop size, set to 1/8th of the window size), WINDOW=“hann” (Hann window function), FMIN=0, and FMAX=24000 (frequency range in Hz). To handle varying call durations, zero-padding was applied to match the length of the longest call, ensuring consistent input dimensions for the classifiers. Each spectrogram was also z-transformed to normalize intensity values across calls, minimizing the effects of amplitude variation. These parameter settings, along with zero-padding and z-transformation, were all based on the suggestions of previous study (Thomas et al., 2022). This process resulted in 10,360 dimensions for each call.

#### Data structure analysis with UMAP

To assess the separability of individual classes and age classes, we performed dimensionality reduction using Uniform Manifold Approximation and Projection (UMAP) from *umap-learn* (McInnes et al., 2018). We repeated supervised UMAP and unsupervised UMAP 100 times each using all calls (651 calls). Supervised UMAP aims to create a latent space where labels (e.g., individual identity) help guide the projection, making it easier to see how well the classes separate. In contrast, unsupervised UMAP does not use labels and reveals the natural structure of the data. In this study, we used two types of labels: individual and age-class (older (over 20 years) and younger (less than 10 years)). Both methods used the same parameters as those in the previous research (Thomas et al., 2022) except for the n_components used for comparison in the previous research: n_components=2, metric=’euclidean’; min_dist=0. We calculated silhouette scores and distances among calls using each UMAP projection to evaluate cohesion within the same class and separation between different classes. Silhouette scores range from -1 to 1, indicating how tightly data points are grouped within a cluster and how well clusters are separated here; higher positive values mean better separation, while higher negative values indicate overlap between clusters.

#### Train-test split

Given the unbalanced sample sizes among individuals, we randomly selected 80 coo calls per individual in each classifier learning iteration. As with the 80 calls in each individual, we allocated 64 calls to the training data and the remainder (16 calls) to the test data. This process was repeated 1000 times, resulting in 1000 pairs of training and test datasets to account for variability and provide a robust estimate of classifier performance.

#### Classifier learning

In this study, we tackled two tasks: One was to identify individuals from the calls of six adult females, and the other was to classify age classes (older = over 20 years old and younger = less than 10 years old). For individual and age-class classification, we trained random forests (RF) and support vector machines (SVM) using training datasets and then predicted test datasets 1000 times using *scikit-learn,* a commonly used Python module (Pedregosa et al., 2011). In each iteration, to estimate the performance of classifiers, we calculated balanced accuracy, which is defined as the average of recall in each class and reduced the biases due to the differences in data size between classes compared to the standard accuracy. By using the prediction based on the test data, we calculated the accuracy per call defined as follows: the number of times the predicted label in the test data matched the actual label divided by the number of times the call was used as the test data.

#### Hyperparameter tuning

Since it is time-consuming to tune the hyperparameters of the classifiers in each iteration, we explored the best parameters using 5-repeated 5-fold cross-validation in the grid search method. In other words, we repeated the random selection of 80 calls from each individual and 5-fold cross-validation 5 times, and then we searched for the combinations of hyperparameters with the highest average balanced accuracy, commonly known as the grid search method. We searched for the best combination of hyperparameters with high average balanced accuracy for RF and SVM in the age-class classification and individual identification tasks, respectively. The grids explored by RF and SVM are as follows: for RF i) max_depth (10, 20, 50, None), ii) max_features (sqrt, log2, None), iii) criterion (gini, entropy, log_loss), iv) min_samples_split (2, 4, 6, 8), v) min_samples_leaf (1, 2, 5); for SVM i) the kernel type (linear, poly, rbf, sigmoid), ii) C, which is the regularization parameter (0.1, 1, 10, 100), iii) the kernel coefficient in poly, rbf and sigmoid (scale, 0.001, 0.01, 0.1, 1), iv) decision_function_shape (ovo, ovr), v) for poly or sigmoid, coef0 (0.0, 0.1, 0.5, 1), and vi) for poly, degree (1, 2, 3, 4). To reduce the computational costs, we set n_estimator =100 in the RF. The best hyperparameters were as follows: In the individual identification task, for RF best hyperparameter combination was n_estimators=100, max_depth=20, max_features=sqrt, criterion=entropy, min_samples_split=2, and min_samples_leaf=1; for SVM C=10, kernel=rbf, gamma=scale, and decision_function_shape=ovo. In the age-class classification task, for RF the best hyperparameter combination was n_estimators=100, max_depth=20, max_features=sqrt, criterion=gini, min_samples_split=2, and min_samples_leaf=2; for SVM C=0.1, kernel=poly, degree=3, coef0=0.5, gamma=0.001, and decision_function_shape=ovo.

## Results

### Individual identification

Coo calls were generally clustered by individual in the supervised UMAP, while the unsupervised UMAP did not produce clear clusters (Fig. 1). This was also reflected in the results of the silhouette score. The mean silhouette score for all six individuals of the supervised UMAP (0.60) was much higher compared to the unsupervised UMAP (0.05) (Fig. 2). The mean silhouette score in each individual was positive (0.21-0.90) in the supervised UMAP; however, the silhouette scores of Kapa and Rine were negative in the unsupervised UMAP. The all-average distances within individuals were smaller than between individuals in the supervised UMAP; on the other hand, the unsupervised UMAP did not show much difference in the within-individual and between-individual distances (Fig. 2).

**Fig. 1.**
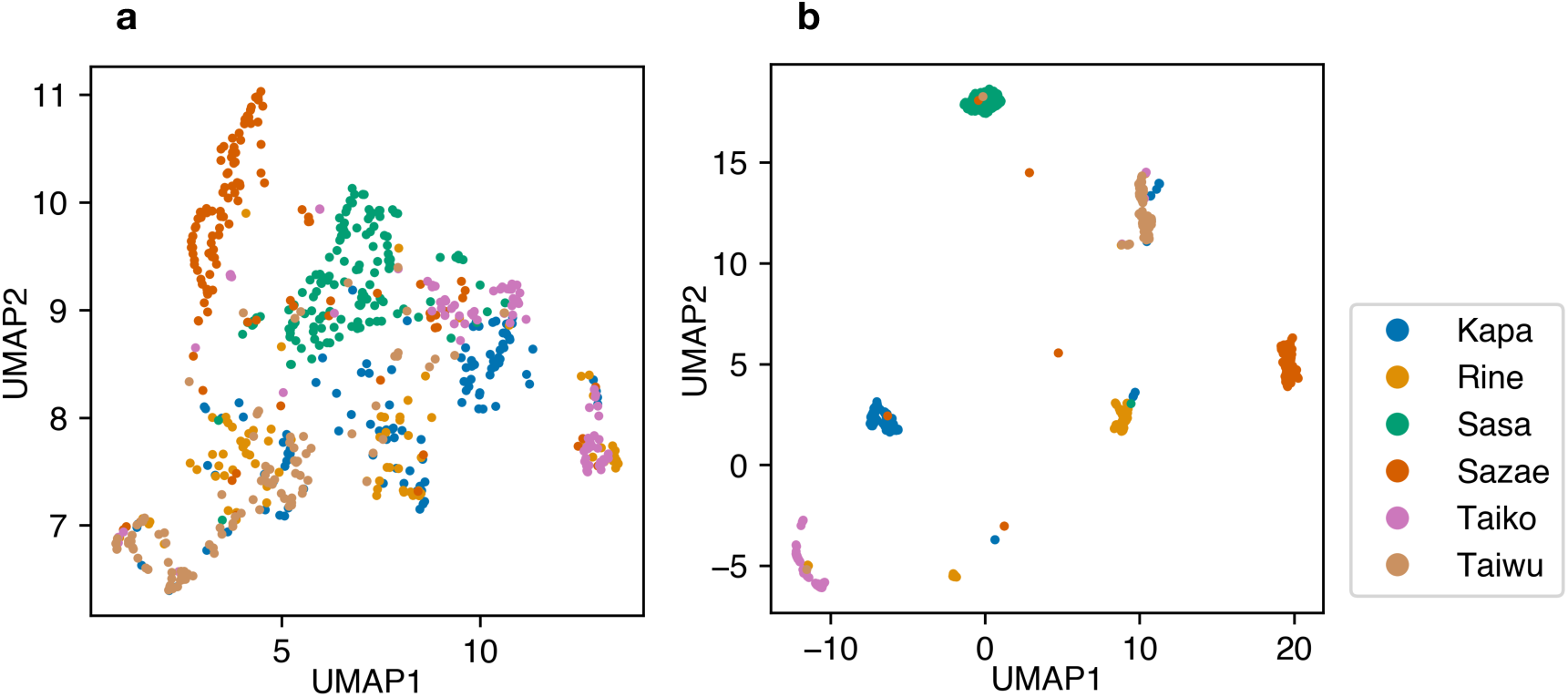
Projections of all coo calls into two-dimensional space through UMAP using individual labels (651 calls; each dot = 1 call; different colors indicate different individuals). a: unsupervised UMAP. b: supervised UMAP.

**Fig. 2.**
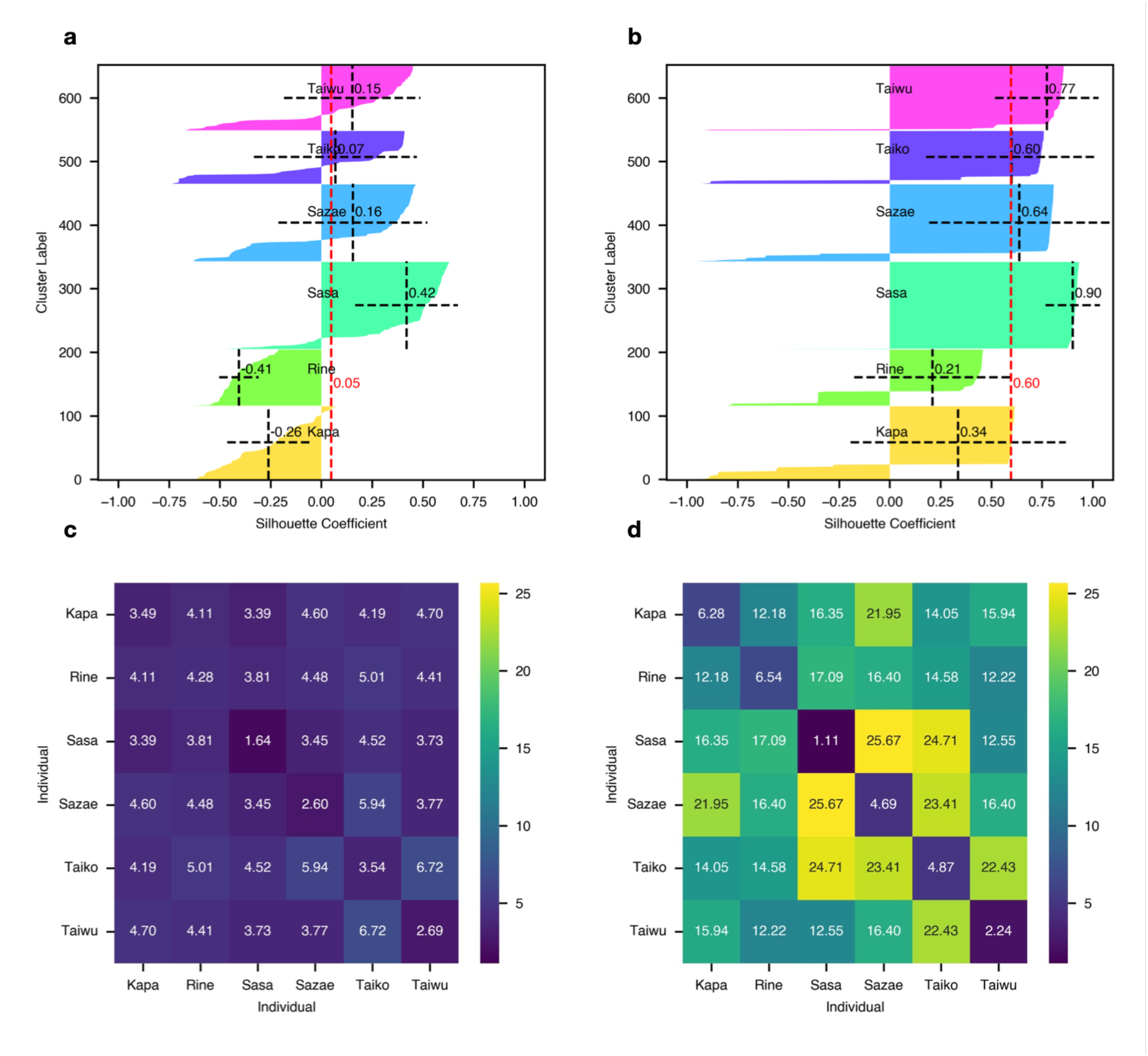
Silhouette plots and distances between individuals using projections of UMAPs and individual labels. Top: silhouette plots of unsupervised (a) and supervised (b) UMAP with averages and standard deviations per individual (black line) and in all individuals (red line). Bottom: distance matrix of unsupervised (c) and supervised (d) UMAP.

**Fig. 3.**
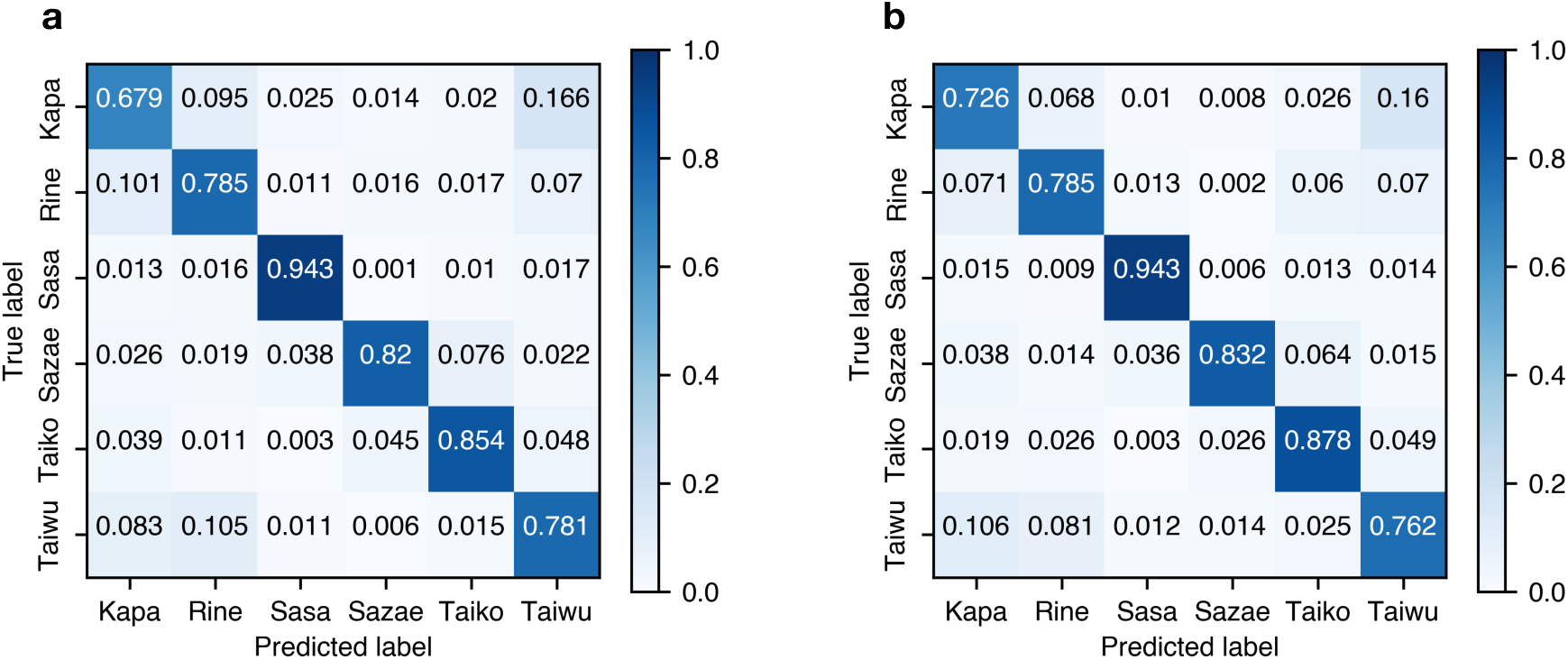
Average confusion matrix for 1000 iterations in identifying six females. a: RF. b: SVM.

In the individual identification task, the mean balanced accuracy of RF was 81% and that of SVM was 82% (Table 1). The mean accuracy varied across individuals, with values ranging from 0.68 to 0.94 in RF and from 0.72 to 0.94 in SVM. Both RF and SVM had the highest accuracy in Sasa and the lowest in Kapa. The main factor lowering the accuracy for both RF and SVM was the misidentification of Kapa, Rine, and Taiwu among them.

**Table 1.** Balanced accuracy of six individual identifications.

| Classifier | Mean | std | min | 25% | 50% | 75% | max |
| --- | --- | --- | --- | --- | --- | --- | --- |
| **RF** | **0.81** | **0.04** | **0.69** | **0.78** | **0.81** | **0.83** | **0.92** |
| **SVM** | **0.82** | **0.04** | **0.71** | **0.79** | **0.82** | **0.84** | **0.93** |

The distribution of the accuracies of all coo calls was divided into high-accuracy and low accuracy calls (Fig. 4). When we defined the calls with accuracy below 1/6 (lower than expected) as low-accuracy calls and over 0.9 as high-accuracy calls, 55 coo calls were classified as low accuracy (Table 2) and 447 as high accuracy (Table 3) in both RF and SVM. The proportion of low-accuracy calls in each individual varied, with the highest for Kapa at 0.15 and the lowest for Sasa at 0.02. On the other hand, the proportion of high-accuracy calls ranged from 0.54 to 0.89.

**Fig. 4.**
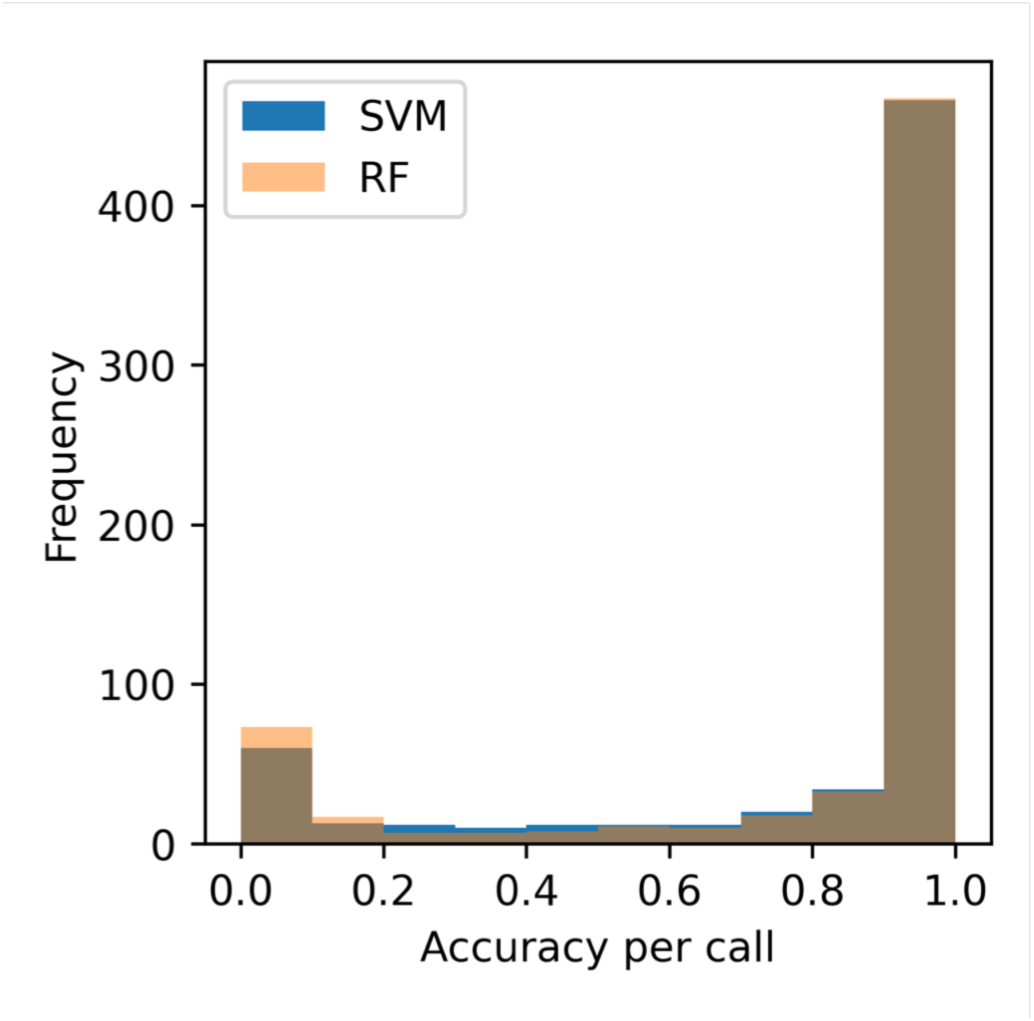
Frequency of accuracy for coo calls in individual identification. Blue bar and orange bar represent SVM results and RF results, respectively.

**Table 2.** Low-and-high accuracy calls in individual identification.

| **Individual** | **Number of**<br>**low-accuracy calls**<br>**in RF** | **Number of**<br>**low-accuracy calls**<br>**in SVM** | **Number of**<br>**low-accuracy calls in**<br>**common between**<br>**RF and SVM** | **Number of calls** |
| --- | --- | --- | --- | --- |
| **Kapa** | **26** | **20** | **17** | **116** |
| **Rine** | **12** | **10** | **9** | **89** |
| **Sasa** | **6** | **6** | **3** | **138** |
| **Sazae** | **18** | **16** | **13** | **122** |
| **Taiko** | **7** | **7** | **4** | **84** |
| **Taiwu** | **12** | **9** | **9** | **102** |

**Table 3.**
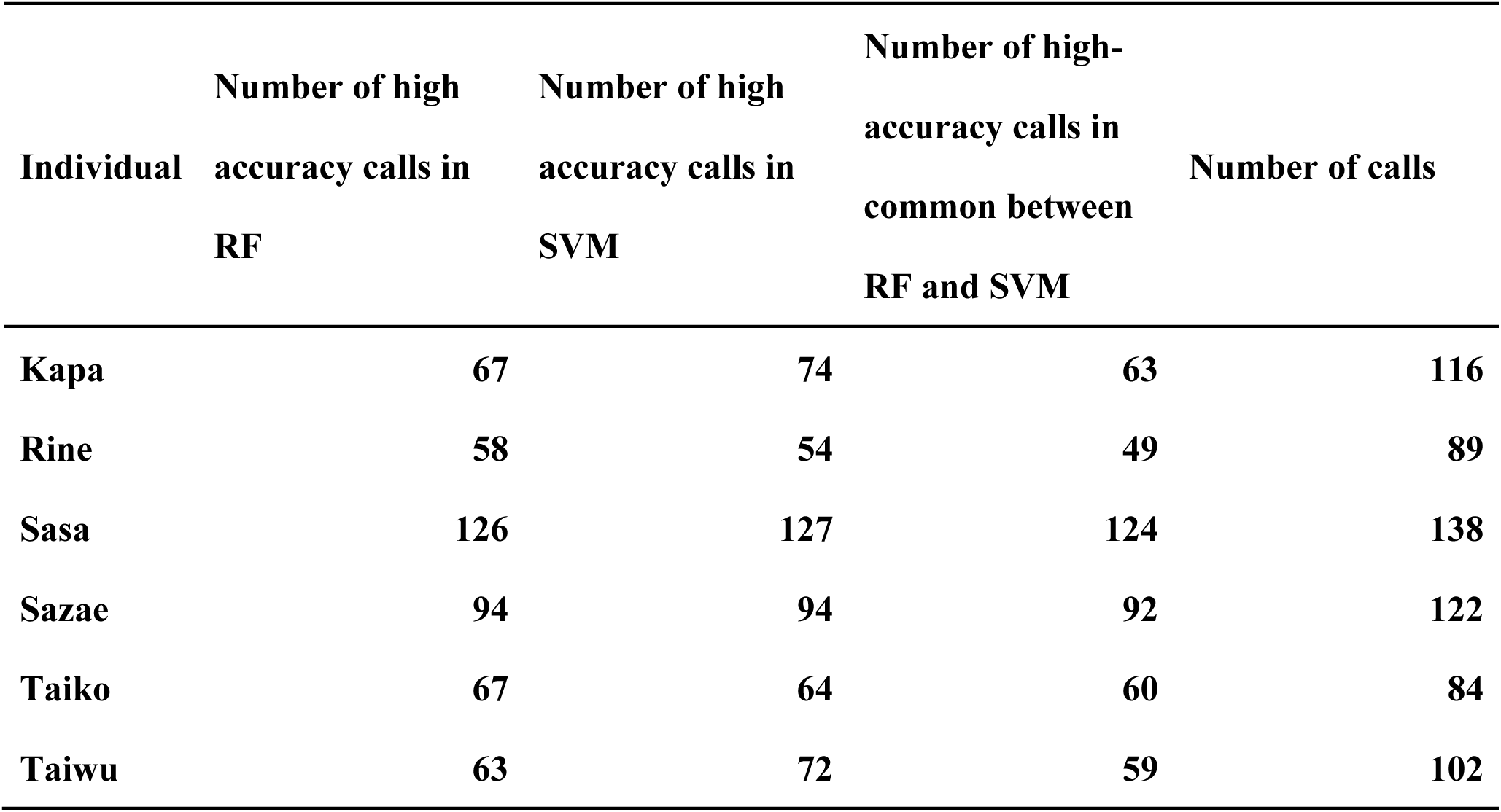
Low-and-high accuracy calls in individual identification.

### Age-class classification

In the supervised UMAP, coo calls were clustered tightly by age-class, while the unsupervised UMAP showed a looser separation between coo calls of younger and older individuals (Fig. 5). As with the supervised UMAP, the averages of silhouette scores were 0.86 for the younger and 0.80 for the older, respectively, which were much higher than the unsupervised UMAP with 0.18 for the younger and 0.01 for the older (Fig. 6). Nevertheless, only some calls of the older individuals had negative silhouette scores in the supervised UMAP. Compared these calls with calls with high silhouette score, but there was no consistent difference between low silhouette calls and high silhouette calls. In terms of distances, the mean pairwise distance within age classes (3.05 for the younger and 3.76 for the older) was smaller than between age classes (22.66) in the supervised UMAP, whereas no differences were observed within and between age classes in the unsupervised UMAP (Fig. 6).

**Fig. 5.**
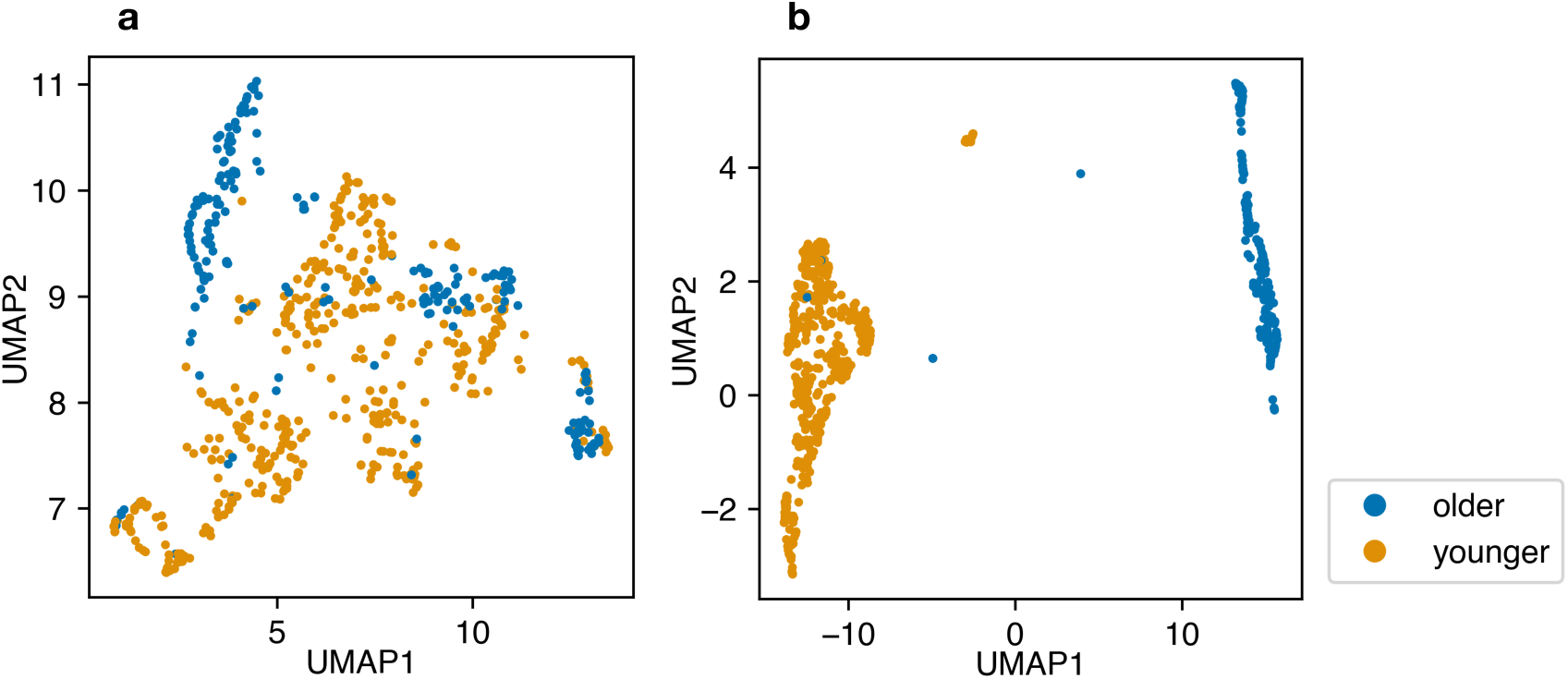
Projections of all coo calls into two-dimensional space through UMAP using age labels (651 calls; each dot = 1 call; different colors indicate different age classes). a: Unsupervised UMAP. b: Supervised UMAP

**Fig. 6.**
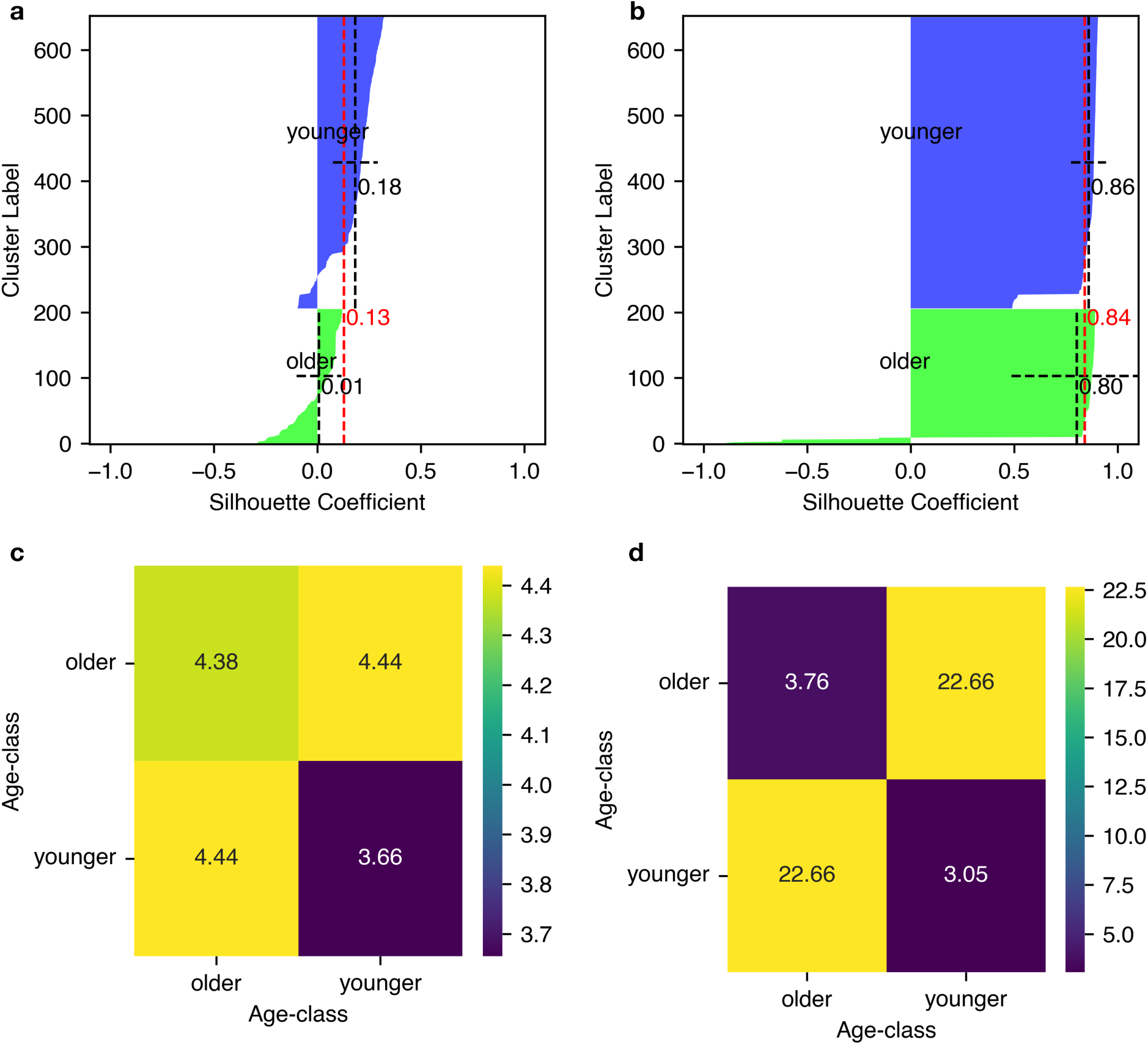
Silhouette plots and distances between individuals using projections of UMAPs and age class labels. Top: silhouette plots of unsupervised (a) and supervised (b) UMAP with averages and standard deviations per age class (black line) and in all classes (red line). Bottom: distance matrix of unsupervised (c) and supervised (d) UMAP.

**Fig. 7.**
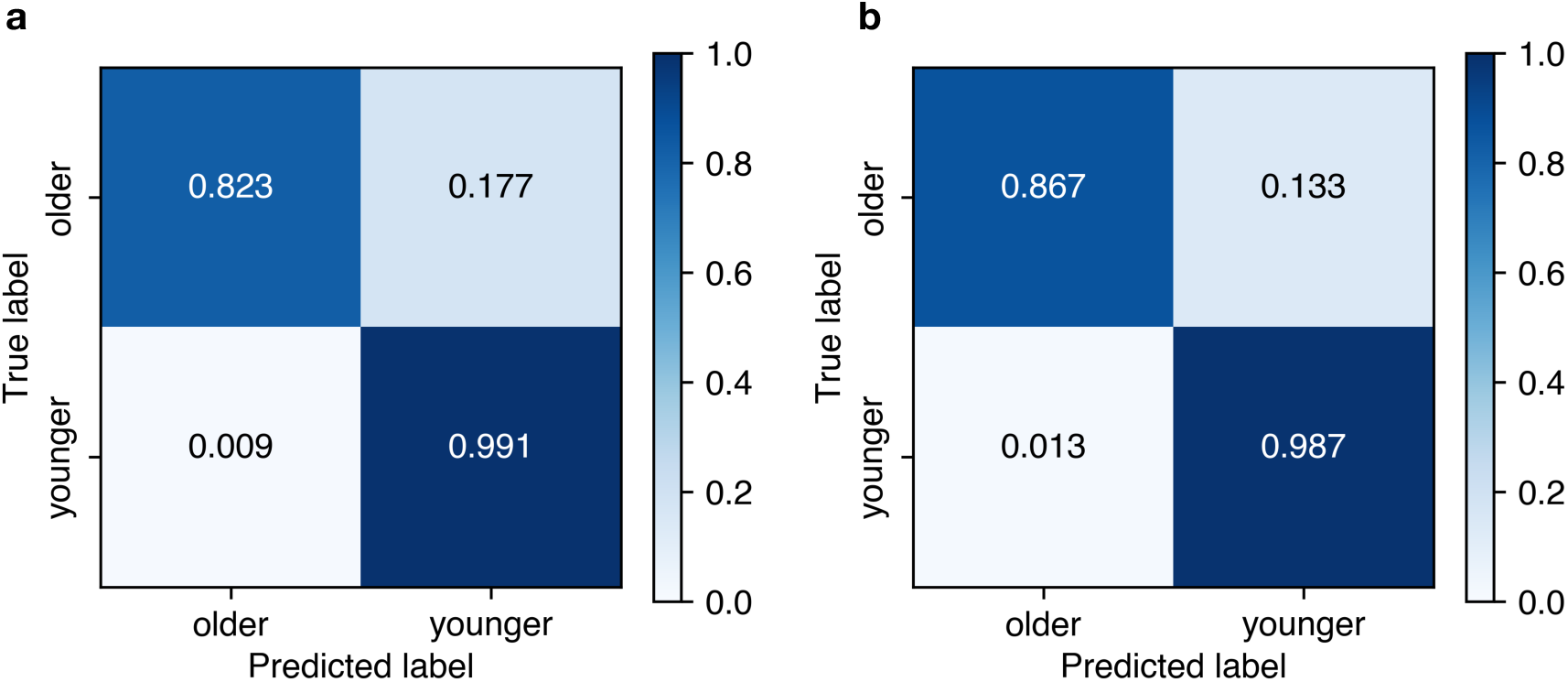
The Confusion matrix for classifying age-class between six females. a: RF. b: SVM.

In the age classification task, the balanced accuracy for RF was 91% and that for SVM was 93% (Table 4), thus exceeding 90%. The accuracy for young individuals exceeded 98% in both RF and SVM, while that for older individuals was less than 87%. Therefore, misclassification was caused by classifying the older calls as younger ones.

**Table 4.** Balanced accuracy of age classification.

| **Classifier** | **Mean** | **std** | **min** | **25%** | **50%** | **75%** | **max** |
| --- | --- | --- | --- | --- | --- | --- | --- |
| **RF** | **0.91** | **0.03** | **0.78** | **0.88** | **0.91** | **0.93** | **0.98** |
| **SVM** | **0.93** | **0.03** | **0.81** | **0.91** | **0.93** | **0.95** | **1.00** |

In the age classification task, the accuracy scores of each call were sharply divided between high and low values (Fig. 8). When we defined the calls with accuracy below 0.5 (lower than expected) as low accuracy calls, the common low-accuracy calls between RF and SVM were 23, comprising of only Sazae and Taiko (Table 5). Here 12.3% of the Sazae calls and 9.5% of Taiko calls were low accuracy.

**Fig. 8.**
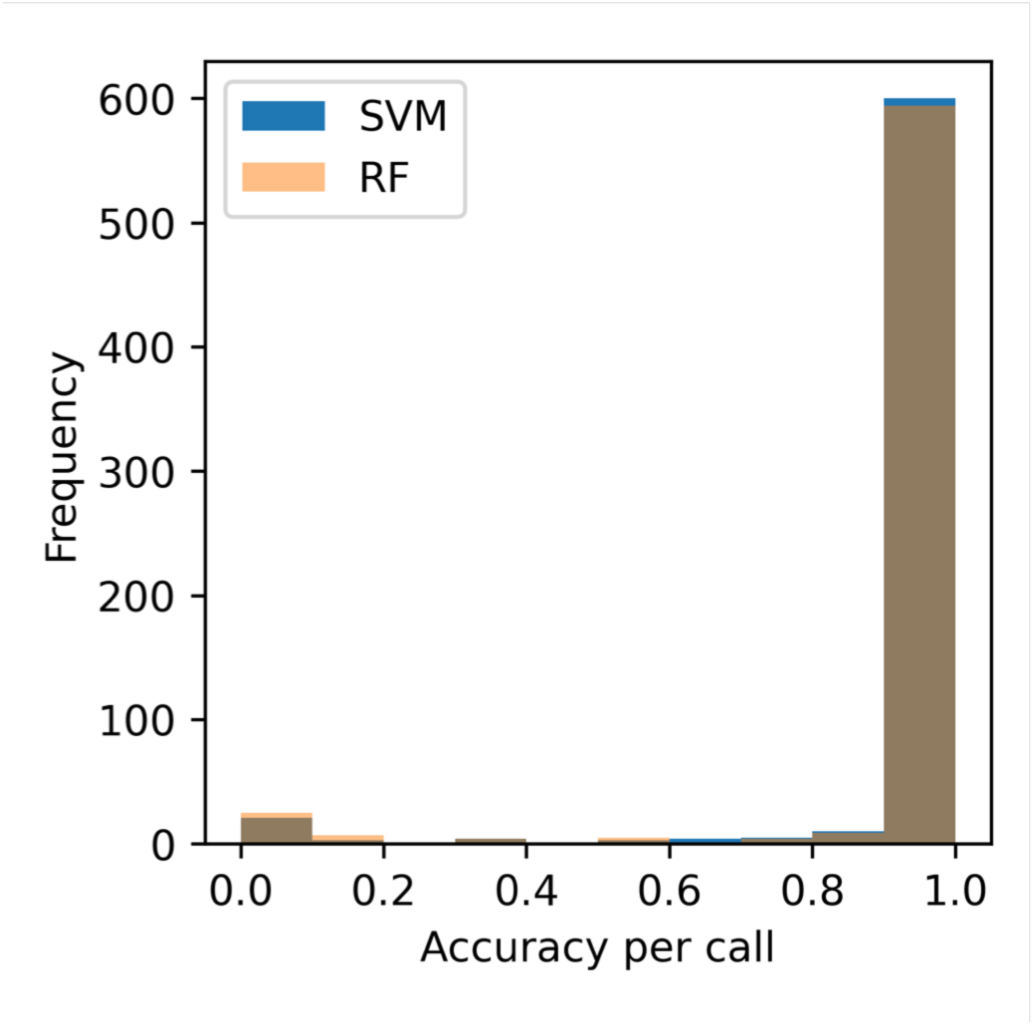
Frequency of accuracy for coo calls in age-class classification. Blue bar and orange bar represent SVM results and RF results, respectively

**Table 5.** Low-accuracy calls in age classification.

| **Individual** | **Number of**<br>**low-accuracy calls**<br>**in RF** | **Number of**<br>**low-accuracy calls**<br>**in SVM** | **Number of**<br>**low-accuracy calls in**<br>**common between RF**<br>**and SVM** | **Number of calls** |
| --- | --- | --- | --- | --- |
| **Kapa** | **0** | **2** | **0** | **116** |
| **Rine** | **0** | **0** | **0** | **89** |
| **Sasa** | **1** | **0** | **0** | **138** |
| **Sazae** | **23** | **16** | **15** | **122** |
| **Taiko** | **14** | **10** | **8** | **84** |
| **Taiwu** | **0** | **1** | **0** | **102** |

## Discussion

### Individual identification

In this study, RF and SVM achieved 81% and 82%, respectively, in identifying the individual from coo calls. The previous research on individual identification from contact calls in macaques achieved variable accuracy (0.32-0.92), although the datasets were obtained in laboratory settings (Ceugniet and Izumi, 2004; Fukushima et al., 2015; Trawicki, 2024). Although it is difficult to simply compare accuracy because of the datasets are different, the dataset used by Fukushima et al. whose study achieved the highest accuracy (0.92) and Trawicki more clustered individually in the UMAP latent space than this study (Sainburg et al., 2020). Therefore, the dataset is probably easier to classify than this study dataset. In addition, considering the difficulty of collecting a balanced set of vocalizations of sufficient and well-controlled sound quality from a social group of wild macaques, the performance of our approach seemed not to be inferior to classifiers that relied solely on highly controlled calls in a laboratory setting. In previous studies, the performance of classifiers varied greatly depending on the acoustic features used (Fukushima et al., 2015; Rendall et al., 1998). This means that the way of selecting acoustic features may distort the performance evaluation. Our approach using mel spectrograms would be more stable due to eliminating the variance of selecting specific acoustic features.

The method using mel spectrograms for individual identification has the potential to facilitate density estimation in the wild. Recently, vocal individual identification has received attention for its use in mark-and-recapture methods due to its non-invasive nature and ability to collect data even at night (Longden et al., 2020). While the average accuracy of individual identification in this study was 80%, more than half of calls achieved over 90% accuracy for any individual with either the RF or SVM models. Highly accurate discrimination is possible by collecting enough data for each individual, identifying calls that are easier to discriminate in advance, and focusing on those calls. Additionally, employing supervised UMAP to pre-check the data structure, as recommended in a previous study (Arnaud et al., 2023), would be beneficial, since supervised UMAP can capture structures that unsupervised UMAP cannot clarify, such as higher silhouette scores and longer inter-class distances (Figs. 1, 2, 5, and 6). Although this study demonstrates the effectiveness of mel spectrograms in small groups, further research is needed to verify whether similar accuracy can be achieved with larger numbers of individuals including not only females but also males with various age-class.

In the individual identification task, the accuracy of each call was sharply divided and skewed into high and the low scores, and low accuracy calls were made by all individuals. These results might be caused by mislabeling or data sampling bias, such as the collection of only a small number of specific types of coo calls in an individual. Otherwise, the results suggest that some of the individuals’ calls might be distinctive. Such differences in individuality might differ depending on the context. For example, in some primates, the call tends to be less individualistic when the distance from other individuals is sufficiently close (Leliveld et al., 2011; Mitani et al., 1996). In addition, the coo call in response tends to resemble the call of the individual whose call was responded to (Sugiura, 1998). These factors may affect the individuality of each coo call. Further study of the relationship between individuality and the context of the calls would help to elucidate which factor affects the difference in individuality.

### Age-class classification

This study achieved 91-93% accuracy in the age-class classification task, i.e. young (<10 yrs) and old (20> yrs) individuals. One possible reason for this high accuracy is the use of the entire mel spectrogram, which preserves global spectral patterns rather than relying on a limited set of predefined acoustic features. However, whether age-class classification based on such representations can be generalized across a broader range of age groups remains to be examined. Because mel spectrograms encode the overall distribution and temporal structure of acoustic energy, they may implicitly reflect complex spectral characteristics. For example, harshness, previously reported to increase with age in Japanese macaques (Inoue, 1988), is difficult to capture using specific acoustic features such as pitch or resonance frequency. Thus, the current results do not demonstrate that the classifiers explicitly relied on harshness or other specific age-related acoustic features, nor do they allow identification of which acoustic properties may have implicitly contributed to the classification.

A key limitation of the present study is that the classifiers may have discriminated age classes based on individual-specific characteristics of a small number of older females rather than generalizable age-related vocal features. In this study, only two older females were included, and although we did not observe an obvious difference in body size between older and younger individuals, we cannot rule out the possibility that subtle individual differences influenced classification. The limited sample size of older individuals therefore represents an important constraint when interpreting the results. Classification accuracy was lower for older individuals than for younger individuals. One possible explanation is that older individuals may not consistently produce vocalizations with age-typical acoustic characteristics, and may, in certain contexts, emit calls that are acoustically similar to those of younger individuals. This variability could reduce classification accuracy for older individuals. However, further investigation is required to evaluate whether such tendencies systematically exist and whether classifiers can robustly capture age-class-specific vocal characteristics.

Taken together, the present findings demonstrate that age-class classification of wild Japanese macaque vocalizations is feasible without explicitly measuring specific acoustic features. At the same time, clarifying which acoustic properties underlie this classification will require future studies incorporating a larger number of older individuals, the use of independent test data from novel individuals, and analytical approaches that allow for greater interpretability of model decisions.

## Acknowledgements

We appreciate the residents and researchers on Yakushima Island for supporting our field study, as well as the Yakushima Forest Ecosystem Conservation Center and the Kagoshima Prefectural Government for permission to perform the research. We also Drs. Hideki Sugiura and Mariko Suzuki for giving thoughtful comments and Sarah Katherine Carnes for preprocessing the vocal data. The study was supported by the Japan Society for the Promotion of Science, Grant-in-Aid for Scientific Research (No. 23H02563 to GH, No. 24H00774 to IM, and No. 23K25168 and No. 23H05428 to HK), JSPS Core-to-Core Program, Asia–Africa Science Platforms (JPJSCCB20250006) and the Collaborative Research Program of Wildlife Research Center, Kyoto University.

## Author contributions

Convinced procedures: RK(lead), FK(supporting), HK(equally), IM (supporting), GH (supporting); Data collection: RK (lead), FK(equally); Software: RK (lead), HK(supporting); Grant obtained: HK(supporting); IM (supporting); GH (lead); Supervision; HK(supporting), IM (supporting), GH (lead); Formal analysis: RK(lead), HK(supporting); Manuscript writing: RK(equally), IM(supporting), HK(lead), GH (supporting)

